# Traumatic Joint Injury Induces Acute Catabolic Bone Turnover Concurrent with Articular Cartilage Damage in a Rat Model of Post-Traumatic Osteoarthritis

**DOI:** 10.1101/2020.04.10.035709

**Authors:** Tristan Maerz, Michael D. Newton, Mackenzie Fleischer, Samantha E. Hartner, Karissa Gawronski, Lucas Junginger, Kevin C. Baker

**Affiliations:** Department of Orthopaedic Surgery, University of Michigan, Ann Arbor, MI; Orthopaedic Research Laboratory, Beaumont Health, Royal Oak, MI; Department of Orthopaedic Surgery, Oakland University William Beaumont School of Medicine, Rochester, MI

## Abstract

**Objective:** Assess acute alterations in bone turnover, microstructure, and histomorphometry following noninvasive anterior cruciate ligament rupture (ACLR).

**Methods:** Twelve female Lewis rats were randomized to receive noninvasive ACLR or Sham loading (n=6/group). I*n vivo* μCT was performed at 3, 7, 10, and 14 days post-injury to quantify compartment-dependent subchondral (SCB) and epiphyseal trabecular bone remodeling. Near-infrared (NIR) molecular imaging was used to measure *in vivo* bone anabolism (800 CW BoneTag) and catabolism (Cat K 680 FAST). Metaphyseal bone remodeling and articular cartilage morphology was quantified using *ex vivo* μCT and contrast-enhanced µCT, respectively. Calcein-based dynamic histomorphometry was used to quantify bone formation. OARSI scoring was used to assess joint degeneration, and osteoclast number was quantified on TRAP stained-sections.

**Results:** ACLR induced acute catabolic bone remodeling in subchondral, epiphyseal, and metaphyseal compartments. Thinning of medial femoral condyle (MFC) SCB was observed as early as 7 days post-injury, while lateral femoral condyles (LFC) exhibited SCB gains. Trabecular thinning was observed in MFC epiphyseal bone, with minimal changes to LFC. NIR imaging demonstrated immediate and sustained reduction of bone anabolism (∼15-20%), and a ∼32% increase in bone catabolism at 14 days, compared to contralateral limbs. These findings were corroborated by reduced bone formation rate and increased osteoclast numbers, observed histologically. ACLR-injured femora had significantly elevated OARSI score, cartilage thickness, and cartilage surface deviation.

**Conclusion:** ACL rupture induces immediate and sustained reduction of bone anabolism and overactivation of bone catabolism, with mild-to-moderate articular cartilage damage at 14 days post-injury.

## INTRODUCTION

Post-traumatic osteoarthritis (PTOA) is a degenerative joint condition known to develop following traumatic joint injuries. Anterior cruciate ligament (ACL) rupture is associated with a particularly high risk for PTOA development – approximately 50% of patients develop PTOA 10-15 years after ACL rupture^1-3^. Surgical ACL reconstruction is a successful treatment to alleviate pain and restore joint stability and function, but despite excellent patient-reported outcomes and a high return-to-sport incidence, reconstruction has not been definitively shown to alter the natural history of PTOA^2, 4, 5^. This observation has led to the hypothesis that acute biological events initiated post-injury play a key role in the onset and progression of PTOA, as opposed to primarily chronic joint instability. These biological events, their respective contribution to PTOA pathogenesis, and the mechanisms that regulate them remain poorly understood.

Crosstalk between articular cartilage (AC) and bone is an important component of healthy joint homeostasis. AC and underlying bone interact via biochemical and mechanical mechanisms, and subchondral bone (SCB) is responsible for mediating the nutritional supply of AC^6^. In both OA and PTOA, the osteochondral unit undergoes dynamic degenerative changes^7, 8^ and increased biochemical crosstalk is hypothesized to drive SCB remodeling, vascular invasion of deep cartilage, and chondrocyte hypertrophy^9-12^. Recent clinical and preclinical studies suggest that alterations in SCB structure and composition may precede detectable damage in AC in idiopathic OA^13-16^, however this has not been shown for PTOA specifically. In PTOA, bony alterations manifest dynamically and are characterized by both lytic and sclerotic phenotypes; SCB loss is observed in the acute timeframe after injury and has been associated with inflammation-mediated osteoclastogenesis and limb offloading due to injury-related gait changes^17, 18^. Chronically, SCB undergoes sclerosis, a classical symptom of radiographic OA recently associated with overactivation of Wnt/β-catenin signaling^19-21^.

Studies of preclinical PTOA have demonstrated rapid loss of SCB and epiphyseal trabecular bone following ACL injury^18, 22-24^. In a murine model of noninvasive ACL rupture-induced PTOA, epiphyseal trabecular bone loss was observed as early as 1-week post-injury, characterized by decreased bone volume fraction, bone mineral density, and trabecular thickness^18, 22^. Using a rat model of noninvasive ACL rupture, our group has demonstrated a similar phenotype of subchondral and epiphyseal trabecular bone loss at intermediate (4-week) and chronic (10-week) timepoints^25^. However, no study has longitudinally characterized bone remodeling *in vivo* acutely after injury, and it remains unclear whether it is associated with dampened bone formation, increased bone resorption, or both. To this end, the purpose of this study was to use *in vivo* imaging, structural histology, and dynamic histomorphometry to characterize acute changes to bone deposition, bone resorption, and bone microstructure in a noninvasive rat model of ACL rupture. Further, we sought to employ quantitative contrast-enhanced μCT and histological evaluation of AC to contextualize bone-related findings with AC degeneration and demonstrate whether acute alterations in bone precede measurable AC changes. We hypothesized that bone anabolism is thwarted and bone resorption (i.e. osteoclast activity) is increased immediately following joint injury, and that major structural changes in SCB and epiphyseal trabecular bone precede marked AC degeneration.

## METHODS

### Animals and Induction of Noninvasive ACL Rupture

Following institutional animal care and use committee approval, twelve female Lewis rats aged 14 weeks, ∼200-220g (Charles River Laboratories, Wilmington, MA, USA) were randomized to ACL rupture (ACLR) or sham injury (Sham) (n=6/group). Sample size was determined based on μCT data from our prior studies using this rat model^25^ based on detection of a 5% difference in trabecular bone volume fraction between groups (effect size=2.2, α=0.05, power=0.9). Immediately prior to injury, rats were administered 5 mg/kg subcutaneous Carprofen, anesthetized with intraperitoneal ketamine/xylazine, and maintained under 1-2% inhaled isoflurane. Noninvasive ACLR was induced using tibial compression-based mechanical loading, as previously described^25-27^. Rats were positioned prone on a custom fixture, with the right knee in ∼100° of flexion. Following preloading (3 N) and preconditioning (1-5 N), a rapid 3.0 mm displacement was applied to the tibia using a mechanical testing system (Insight 5, MTS Systems, Eden Prairie, MN, USA), resulting in a closed, isolated ACL rupture. Sham rats underwent preload and preconditioning only, without 3-mm injury loading. Following loading, animals were administered the anesthetic reversal agent yohimbine (0.2 mg/kg SC). Rats were allowed *ad libidum* cage activity in a 12-hr light/dark facility. To enable dynamic histomorphometric assessment of bone formation, rats received intraperitoneal injections of 1% calcein solution buffered with NaHCO_3_ at the time of injury and 24 hrs prior to CO_2_ asphyxia-induced euthanasia 14 days post-injury.

### Near Infrared (NIR) Molecular Imaging to Assess Bone Deposition and Resorption

At 3, 7, 10, and 14 days post-injury, rats underwent *in vivo* near-infrared fluorescence (NIR) molecular imaging to longitudinally quantify bone deposition. Twenty-four hours prior to imaging, rats received 5 nmol intravenous IRDye 800 CW BoneTag (BoneTag) (LI-COR, Lincoln, NE, USA). BoneTag is a calcium-chelating fluorescent compound that incorporates into newly mineralized bone, enabling *in vivo* assessment of bone formation^28^. At the 14-day timepoint, to assess *in vivo* osteoclast activity as a measure of bone resorption, rats received 5 nmol Cat K 680 FAST (CatK) (PerkinElmer, Waltham, MA, USA). CatK is an activatable fluorescent probe that detects *in vivo* Cathepsin K activity, an indirect measure of osteoclast activity^29, 30^. On the day of imaging, lower limb fur was removed, and lateral NIR images of both hindlimbs were acquired in the 700 and 800 nm channels (Pearl Impulse, LI-COR) under isoflurane-induced anesthesia. Consistently-sized regions of interest (ROIs) were virtually placed onto each knee (Fig 3A), and mean fluorescent intensity was calculated within each ROI. To control for compounding BoneTag signal and animal-to-animal variability, normalized BoneTag and CatK signals were calculated by dividing mean fluorescent intensity of the ACLR injured/Sham uninjured limb by its respective contralateral limb.

**Fig 1.**
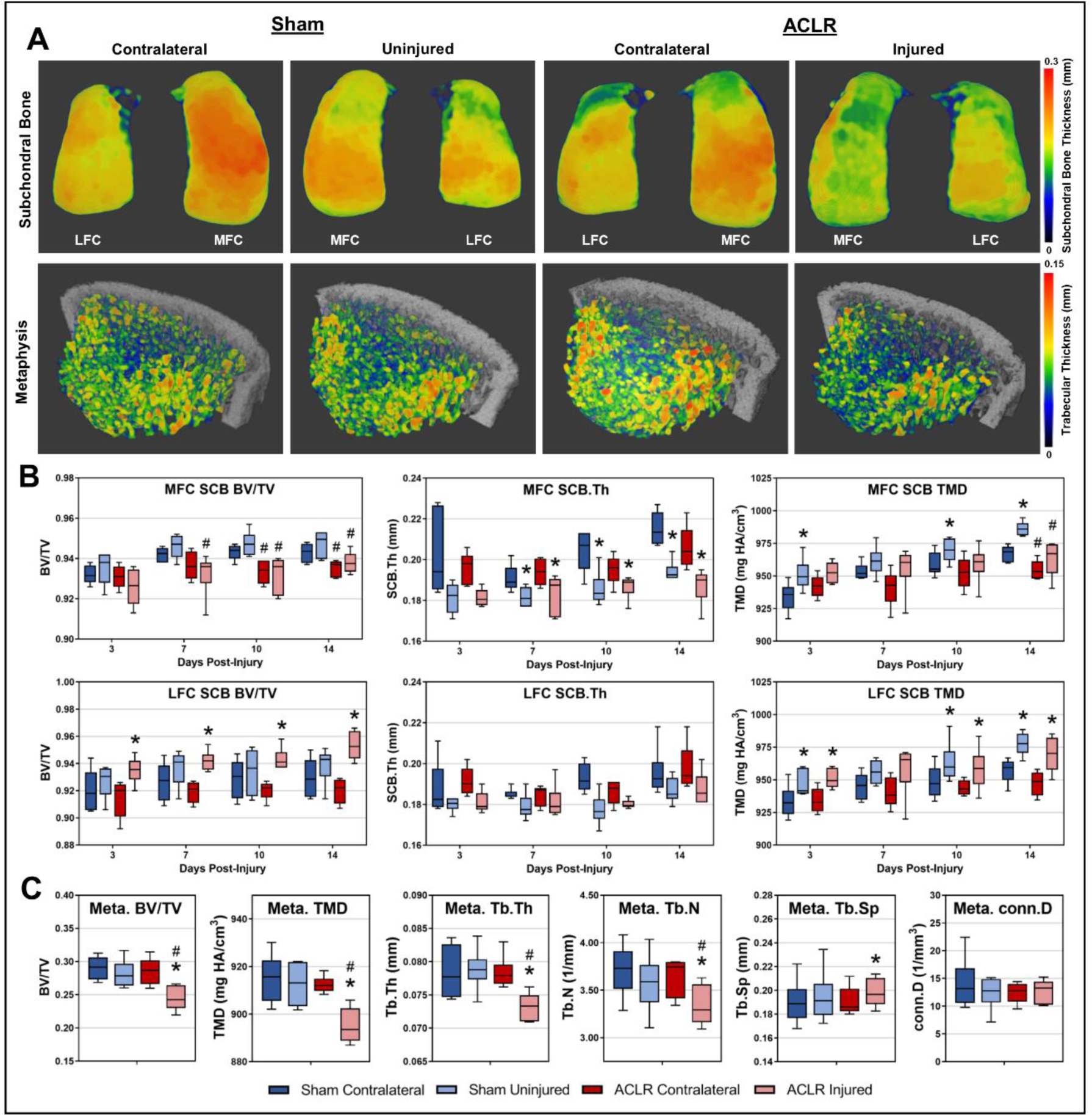
Subchondral and Metaphyseal Bone Morphometry. Three-dimensional thickness maps demonstrate compartment-dependent alterations in SCB morphometry and catabolic remodeling of metaphyseal trabecular bone following ACLR (A). Compared to Sham, ACLR had lower SCB BV/TV and TMD on the MFC in both limbs, whereas the LFC of injured ACLR knees exhibited BV/TV and TMD gains (n=6 Sham; n=5 ACLR) (B). Both ACLR and Sham induced SCB thinning on the MFC, compared to contralateral knees (B). MFC thinning in Sham was confined to the anterior condyle, whereas the ACLR MFC exhibits thinning throughout the entire condyle (B). ACLR also induced catabolic remodeling of metaphyseal trabecular bone (n=6 Sham; n=6 ACLR) (A,C). * indicates significant difference to contralateral limb; # indicates significant difference to ipsilateral limb in Sham.

**Fig 2.**
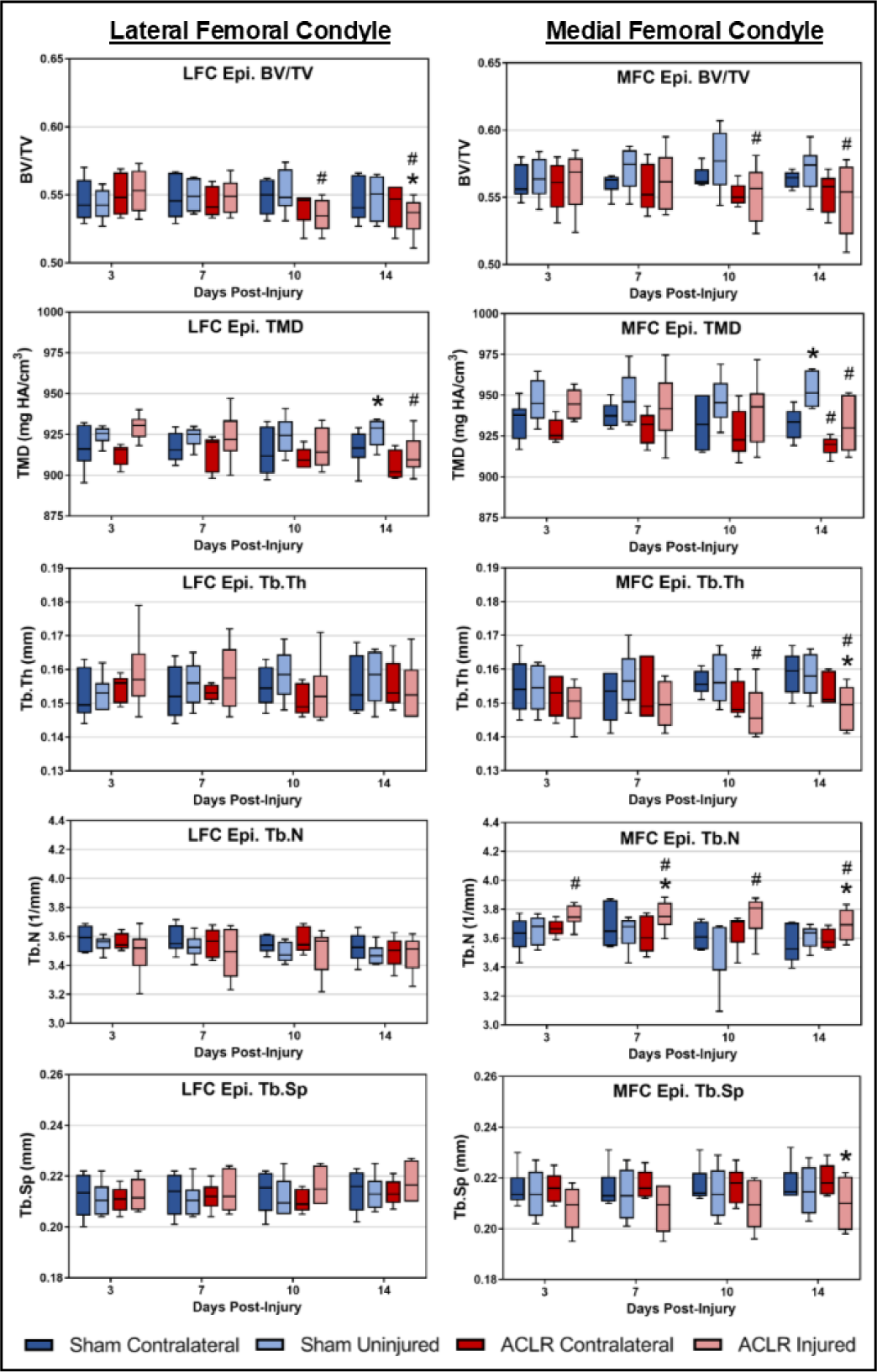
Epiphyseal Bone Morphometry of MFC and LFC. Longitudinal, in vivo µCT of epiphyseal trabecular bone demonstrates that ACLR-induced catabolic remodeling is most pronounced on the MFC. ACLR induces loss of trabecular BV/TV, compared to Sham femora. By 14d post-injury, injured ACLR femora exhibit decreased Tb.Th, decreased Tb.Sp, and increased Tb.N. No changes in Tb.Th, Tb.N, or Tb.Sp were noted in the LFC in either ACLR or Sham. (n=6 Sham; n=5 ACLR) * indicates significant difference to contralateral limb; # indicates significant difference to ipsilateral limb in Sham.

**Fig 3.**
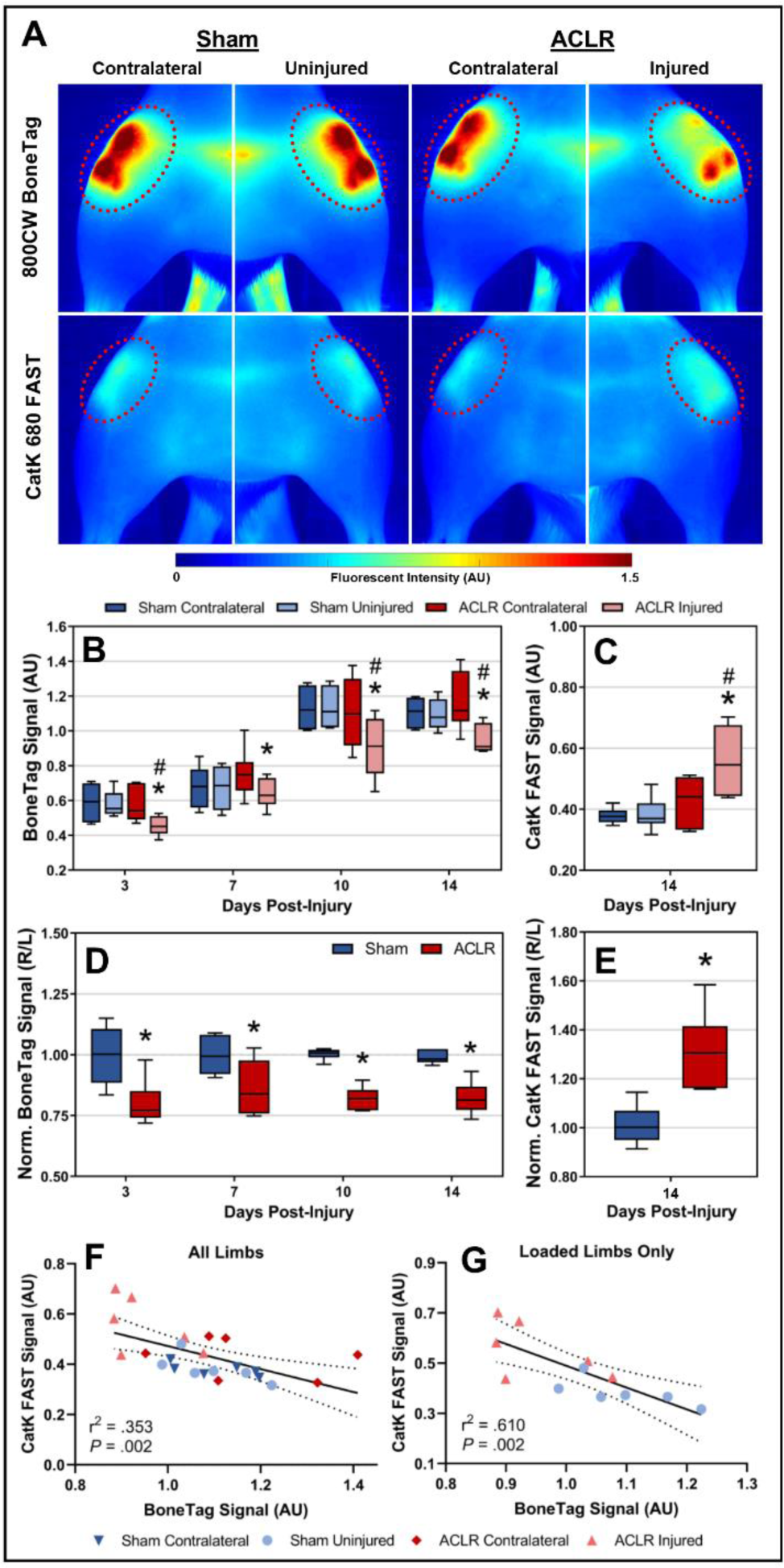
Near-infrared Molecular Imaging of Bone Turnover. NIR heatmaps (A) indicate decreased BoneTag and increased CatK signal in the injured limb of ACLR. Quantitatively, there was a significant reduction in longitudinal BoneTag (B) and significant increase in endpoint CatK (C) signal compared to both Sham ipsilateral and ACLR contralateral limbs (n=6 Sham; n=6 ACLR). Normalized BoneTag (D) and CatK (E) data indicate a ∼15-20% reduction and ∼32% increase, respectively. A poor correlation between BoneTag and CatK signal was observed when all study limbs were analyzed (F), however including only loaded limbs yielded a moderate inverse correlation, demonstrating the in vivo relationship between bone anabolism and catabolism following ACLR (G). * indicates significant difference to contralateral limb; # indicates significant difference to ipsilateral limb in Sham.

### µCT Imaging

Rats underwent live, bilateral *in vivo* µCT imaging of the distal femoral epiphysis at 3, 7, 10, and 14 days post-injury (55 kVp, 114 μA,15.6 μm voxel, Viva-80, Scanco Medical AG, Brüttisellen, Switzerland) under isoflurane-induced anesthesia. Following euthanasia at 14 days, distal femora were harvested and meticulously dissected under microscopy to expose AC, facilitating subsequent contrast-enhanced µCT of AC. Femora were fixed in 10% neutral buffered formalin for 72 hrs and stored in 70% ethanol. At the time of *ex vivo* imaging, femora were rehydrated in PBS for 24 hrs. Distal femoral metaphyses were imaged using *ex vivo* μCT (55 kVp, 145μA, 6 μm voxel, μCT-40, Scanco Medical), as longitudinal *in vivo* imaging was not possible due to live imaging time limitations. Femora were then incubated in 20% ioxaglate (Hexabrix 320, Guerbet LLC, Princeton, NJ), pH=7.2 for 24 hrs, and contrast-enhanced µCT imaging of AC was acquired (55 kVp, 145μA, 8 μm voxel). All *ex vivo* imaging was performed in a humidified sample holder.

### µCT Image Analysis

Image processing and analysis was performed using MATLAB (Mathworks Inc., Natick, MA, USA). All manual tissue contouring was performed by the lead author (TM). Standardized bone morphometry parameters were calculated using the ImageJ plug-in BoneJ^31^, utilizing the MATLAB–ImageJ interface Miji^32^.

To enable accurate voxel-by-voxel characterization of longitudinal bone remodeling, a semi-automated registration algorithm was used to segment epiphyseal trabecular bone and SCB from *in vivo* µCT images. First, epiphyseal trabecular bone and AC were segmented from endpoint contrast-enhanced µCT via manual contouring and automated, registration-based segmentation (described below), respectively. Contrast-enhanced µCT scans for each limb were rigidly registered onto respective *in vivo* µCT data sets, enabling mapping of epiphyseal bone and AC volumes across longitudinal data. SCB was segmented by dilating AC volumes and thresholding bone, as previously described^25^. Metaphyseal trabecular bone was manually contoured from *ex vivo* µCT images. BV/TV, BMD, TMD, Tb.Th, Tb.N, and Tb.Sp were calculated for epiphyseal and metaphyseal trabecular bone volumes, while BV/TV, BMD, TMD, and SCB.Th were calculated for SCB volumes.

AC volumes were segmented from contrast-enhanced µCT images using a custom, atlas-based registration scheme (data not published; manuscript under revision; expected publication June 2020). Briefly, an average tissue atlas of the distal femur was generated from a population of manually segmented training images. The averaging process yields an atlas with pre-defined tissue boundaries and averaged anatomy, enabling robust and highly accurate registration onto both healthy and injured femora (Dice Similarity Coefficient > 0.95). This atlas was registered onto contrast-enhanced µCT images, followed by thresholding to remove residual air and bone, to segment AC. Registrations were inspected by an expert (MDN) to confirm accurate segmentation. AC morphology was analyzed by mean cartilage thickness (MCT) and surface deviation (S_a_), as previously shown by our group^26, 33^. In brief, the bone-cartilage interface was isolated from final AC volumes and mapped from 3D to 2D using conformal parameterization^34^, enabling the generation of 2D AC thickness maps, from which MCT and S_a_ can be derived.

### Structural Histology and Dynamic Histomorphometry

Fixed femora were processed for undecalcified histology and embedded in polymethyl methacrylate. Spaced 6-μm sagittal sections of the medial femoral condyle (MFC) and lateral femoral condyle (LFC) were cut and stained with Safranin-O/Fast Green (Saf-O). Adjacent unstained sections at each interval were cut and mounted for fluorescent imaging of calcein to quantify bone formation. Further sections at each interval were stained for tartrate-resistant acid phosphatase (TRAP) to identify osteoclasts. Brightfield imaging was performed at 20x using an automatic slide imaging system (Aperio, Leica 122 Biosystems, Buffalo Grove, IL, USA). Fluorescent imaging of calcein labeling was performed using standard FITC fluorescent microscopy at 10x magnification (Eclipse E800, Nikon, Tokyo, Japan).

To assess osteoarthritis severity, Saf-O stained sections were evaluated by two blinded investigators using the Osteoarthritis Research Society International (OARSI) score^35^. OARSI scores of the MFC and LFC were evaluated both separately and together as an average score. Static and dynamic histomorphometric parameters were quantified in a blinded fashion using the Bioquant Osteo software (Bioquant Image Analysis Corp., Nashville, TN), according to standard procedures^36^. In brief, Saf-O sections were used to quantify static measures (BV, TV, and BS) in epiphyseal trabecular bone. Adjacent unstained fluorescent sections were used to measure calcein labels to derive dynamic measures of bone formation within the 14-day study period, namely total mineralizing surface (MS), mineralizing surface over bone surface (MS/BS), mineral apposition rate (MAR), bone formation rate over bone volume (BFR/BV). An important distinction in these parameters is that MAR and BFR/BV are measures of bone formation kinetics (directly related to the inter-label distance of calcein bands), whereas MS/BS assesses the proportion of actively mineralizing bone independent of bone formation rates (independent of inter-label distance). Lastly, TRAP^+^ osteoclasts were segmented from TRAP-stained sections using image analysis, and osteoclast number per total area (N.Oc/T.Ar) and osteoclast number per bone surface (N.Oc/BS) were quantified.

### Data Analysis and Statistics

Statistical analyses were performed in SPSS (v22, IBM, Armonk, NY). Normality and equal variance assumptions were confirmed and appropriately addressed in all continuous data. Longitudinal *in vivo* data was analyzed between Sham and ACLR groups and between injured/uninjured and contralateral limbs using a linear mixed model, with “group” as a between-subject factor and “limb/laterality” and “time” as within-subject factors. This analysis enabled us to sensitively elucidate longitudinal trends by appropriately accounting for within-subject and between-subject variance. *Ex vivo*/endpoint µCT data was analyzed by two-way analysis of variance (ANOVA), with “group” as a between-subject factor and “limb/laterality” as a within-subject factor. Multiple comparisons were performed with Sidák P-value correction. Ordinal data was compared using Kruskal-Wallis tests with a Dunn’s *post-hoc* comparison correction. Bivariate correlations were performed using Pearson correlations. Adjusted P-values less than 0.05 were considered significant.

## RESULTS

### ACL Rupture Induced Progressive and Compartment-Dependent Subchondral Bone Alterations

Aging-related gradual increases in SCB BV/TV, TMD, and SCB.Th are evident in the Sham group (Fig 1B). ACLR induced longitudinal loss of SCB BV/TV and SCB.Th (Fig 1B), and, unexpectedly, sham-loaded rats also exhibited significantly lower SCB.Th compared to their respective contralateral femora, albeit to a lesser degree than ACLR (Fig 1B). SCB thinning of the MFC in Sham uninjured limbs was confined to the anterior condyle, whereas the MFC in injured ACLR limbs exhibited thinning through the entire condyle (Fig 1A). By the 14d timepoint, injured knees in ACLR had significantly lower SCB BV/TV and TMD compared to uninjured knees in Sham. On the LFC, injured ACLR femora exhibited progressive SCB BV/TV gains compared to contralateral knees, and loaded knees in both Sham and ACLR exhibited SCB TMD gains (Fig 1B).

### ACL Rupture Induced Catabolic Remodeling of Femoral Metaphyseal and Epiphyseal Trabecular Bone

Compared to both contralateral ACLR femora and uninjured Sham femora, injured ACLR femora exhibited significantly lower metaphyseal BV/TV, TMD, BMD, and Tb.Th. (Fig 1) at the 14d endpoint. Injured ACLR femora also had significantly lower metaphyseal Tb.N and significantly higher Tb.Sp compared to uninjured contralateral femora.

Longitudinal µCT imaging of epiphyseal bone revealed a progressively catabolic phenotype in the MFC and LFC following ACLR, with the most pronounced changes in the MFC (Fig 2). Significant decreases in trabecular BV/TV and Tb.Th of the injured ACLR MFC were observed by the 10d and 14d timepoints (Fig 2), and loss of BV/TV was also observed on the LFC as early as 10d. Both femora in ACLR exhibited decreased epiphyseal TMD compared to respective limbs in Sham, likely due to reduced activity in ACLR rats. Injured ACLR femora exhibited significantly, albeit marginally increased Tb.N in the MFC. No changes in Tb.Th, Tb.N, or Tb.Sp were noted in the LFC in any femora in ACLR or Sham (Fig 2).

### In vivo NIR molecular imaging reveals reduced bone formation and greater catabolic bone turnover following ACLR

To elucidate whether joint injury-induced alterations in bone microstructure are due to dampened anabolic bone formation, due to increased catabolic osteoclast activity, or both, we employed *in vivo* NIR molecular imaging. BoneTag NIR signal – a direct measure of new bone formation – was significantly decreased in ACLR injured limbs at all timepoints compared to contralateral limbs and at 3d, 10d, and 14d compared to Sham uninjured limbs (Fig 3A, B), indicating an immediate and sustained impact of ACLR on bone anabolism. NIR heatmaps demonstrate marked loss of BoneTag signal in ACLR, most notably at the location of the femoral epiphysis, whereas Sham knees exhibit higher, evenly-distributed BoneTag signal throughout the whole joint (Fig 3A). CatK NIR signal – an indirect measure of osteoclast activity^29, 30^ – was measured at the final 14d timepoint and found to be significantly increased in ACLR injured limbs compared to both contralateral ACLR limbs and uninjured Sham limbs (Fig 3C). CatK heatmaps demonstrate increased signal intensity throughout the entire knee joint in injured ACLR limbs (Fig 3A). To control for compounding BoneTag signal and potential contralateral effects (i.e. reduced activity in ACLR rats), we normalized BoneTag and Cat K signal of the injured/uninjured limbs in ACLR/Sham to their respective contralateral limb. Normalized data indicates a sustained ∼15-20% reduction in longitudinal BoneTag signal (Fig 3D) and a ∼32% increase in CatK signal (Fig 3E) in ACLR, whereas normalized BoneTag and CatK signal in Sham were consistently ∼1.

To ascertain the relationship between bone anabolism and bone catabolism, we performed bivariate correlations between absolute CatK and BoneTag signal. Analyzing all limbs of both Sham and ACLR yielded a weak, albeit significant inverse correlation (r^2^=0.353, *P*=0.002) between CatK and BoneTag signal (Fig 3F). Including only loaded limbs (ACLR injured and Sham uninjured) markedly improved this correlation (r^2^=0.610, *P*=0.002) (Fig 3G), indicating an *in vivo* relationship between bone anabolism and catabolism following joint loading in our model. Collectively, this data demonstrates that ACLR is associated with both decreased bone formation (i.e. anabolism) and increased osteoclast activity (i.e. catabolism) and that these processes are inversely correlated *in vivo*.

### Histomorphometry Indicates Thwarted Bone Formation and Increased Osteoclast Number

To label newly-deposited bone and quantitatively assess 14d bone formation, we injected rats with calcein at the beginning and end of the study. Fluorescent microscopy demonstrates reduced overall calcein uptake, a reduced incidence of double calcein labels, and reduced inter-label distance in the epiphyseal trabecular bone of ACLR injured femora (Fig 4A), indicating thwarted bone formation. Uninjured contralateral ACLR femora and both femora of Sham exhibit consistent presence of double labels throughout the epiphysis, indicative of new bone formation. Quantitative measurement of dynamic histomorphometric parameters corroborate the observation of reduced bone formation, and injured ACLR femora had significantly lower epiphyseal MS compared to contralateral limbs and significantly lower BFR/BV compared to uninjured, Sham limbs (Fig 4B). There were no significant differences in epiphyseal MS/BS and MAR (Fig 4B).

**Fig 4.**
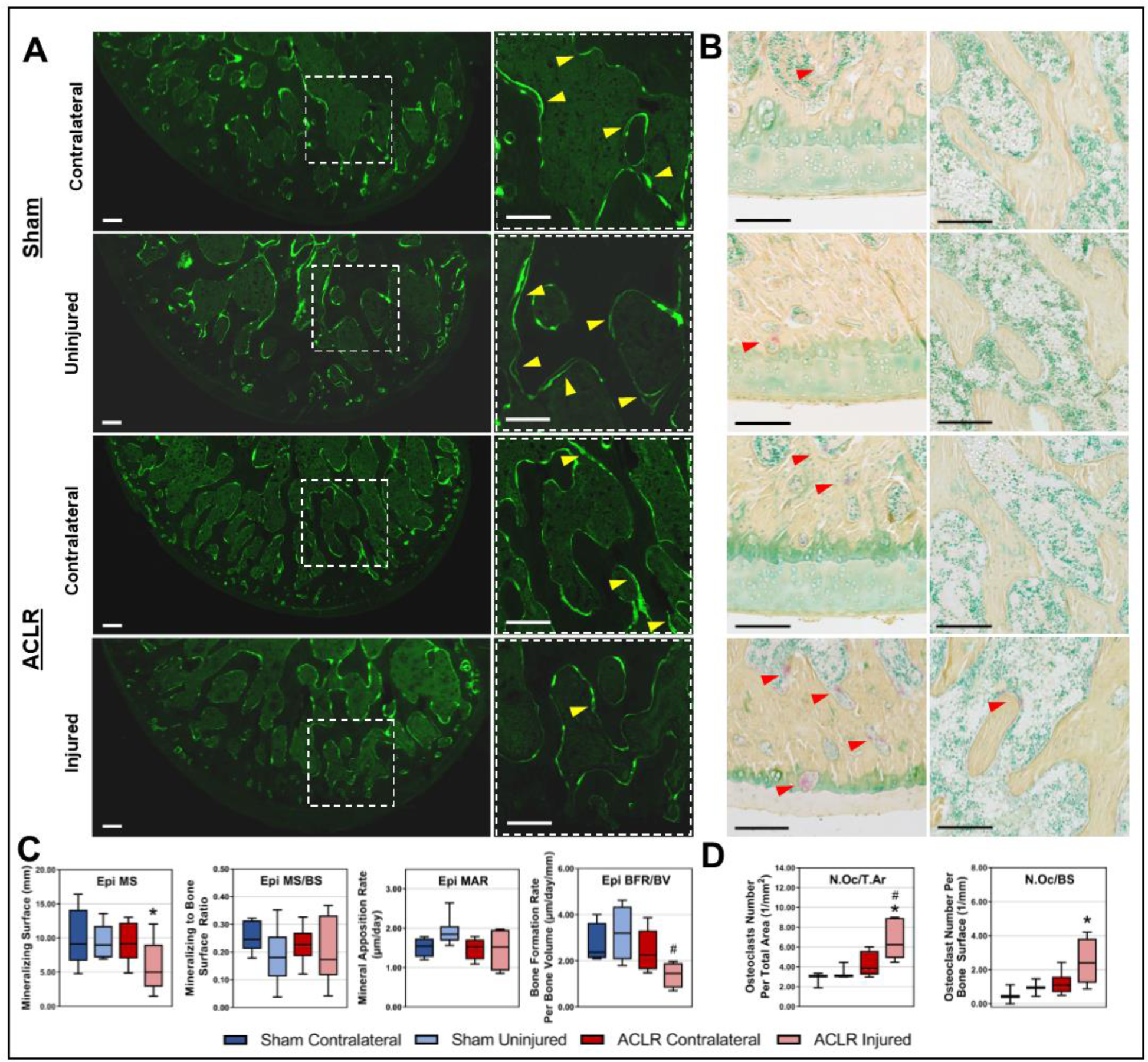
Histomorphometry of Bone Formation and Osteoclast Density. Fluorescent sections (A) demonstrate consistent presence of double calcein labels (yellow arrowheads) in both Sham limbs and in the ACLR contralateral limb. The ACLR injured limb exhibits an overall reduced calcein uptake and a lower incidence of double labels, indicating thwarted bone formation. Quantitatively, ACLR Injured limbs exhibit reductions in epiphyseal MS and BFR/BV (n=5 Sham; n=6 ACLR) (C). Brightfield imaging of TRAP-stained sections indicate increased osteoclast numbers in ACLR Injured femora (B). Osteoclasts (red arrowheads) were most numerous in subchondral bone (B, left column) compared to epiphyseal trabecular bone (B, right column). Quantitatively, ACLR Injured femora had significantly elevated N.Oc/T.Ar and N.Oc/BS (n=3 Sham; n=6 ACLR) (D). * indicates significant difference to contralateral limb; # indicates significant difference to ipsilateral limb in Sham. Calcein images scalebar is 250 μm. TRAP images scalebar is 150 μm.

TRAP-stained slides were analyzed to quantify osteoclast density within the distal femoral epiphysis. Qualitatively, all groups exhibited greater osteoclast numbers in SCB (Fig 4C, left column) compared to trabecular bone (Fig 4C, right column). Injured ACLR femora had more abundant osteoclasts, most notably in or near SCB (Fig 4C, left column) – osteoclasts were commonly observed penetrating calcified cartilage and, in some instances, spanning the tidemark (Fig 4C). Quantitatively, ACLR injured femora had significantly higher N.Oc/T.Ar compared to both ACLR Contralateral and Sham Uninjured and significantly higher N.Oc/BS compared to ACLR Contralateral (Fig 4D). Taken together, histomorphometric assessments demonstrate that ACLR thwarts new bone formation and induces osteoclast infiltration in epiphyseal bone, most notably in the subchondral compartment.

### Early Articular Cartilage Damage is Primarily Confined to the MFC following ACLR

To contextualize bone-related findings with the degree/severity of cartilage degeneration at the acute 14d timepoint after ACL injury, we analyzed femoral AC using contrast-enhanced μCT and sagittal Saf-O-stained histology. Histologically, femoral cartilage of both Sham limbs appeared normal with no alterations in structure, cellular phenotype, or staining intensity (Fig 5A). Minor cartilage damage, most commonly superficial fibrillation, was observed on the MFC of ACLR contralateral femora (Fig 5A). The MFC of ACLR injured femora exhibited marked cartilage damage and degeneration, with the anterior condyle exhibiting structure damage, cartilage erosion, hypocellularity, and loss of Saf-O staining. The posterior condyle exhibited notable swelling, increased chondrocyte cloning and clustering, and damage to the superficial zone (Fig 5A). Compared to both ACLR contralateral and Sham ipsilateral, ACLR injured femora had significantly greater MFC and whole-femur OARSI score, with marginal but non-significant increases in OARSI score on the LFC (Fig 5C). Contrast-enhanced μCT thickness maps corroborate histological findings: Sham femora exhibit a generally congruent articular surface with normal zones of increased cartilage thickness at the weight-bearing regions of the condyles (Fig 5B). Injured ACLR femora exhibit abnormal cartilage thickness distributions, most notably on the MFC, where zones of markedly-increased cartilage thickness were observed at the central weight-bearing region and zones of markedly-decreased cartilage thickness were observed at the anterior aspect (Fig 5B). Quantitatively, both condyles of injured ACLR limbs exhibited significant increases in mean cartilage thickness of ∼20-30% compared to uninjured contralateral and Sham limbs (Fig 5D). Lastly, injured ACLR femora exhibited a significant ∼75-95% increase in MFC S_a_ and a significant ∼50-60% increase in whole-femur S_a_ (Fig 5E). Due to only cartilage thickening and no observed cartilage loss on the LFC of ACLR injured femora, there was no significant increase in LFC S_a_ (Fig 5E). Both Sham femora and the ACLR contralateral femora exhibited similar cartilage thickness and S_a_ (Fig 5D,E).

**Fig 5.**
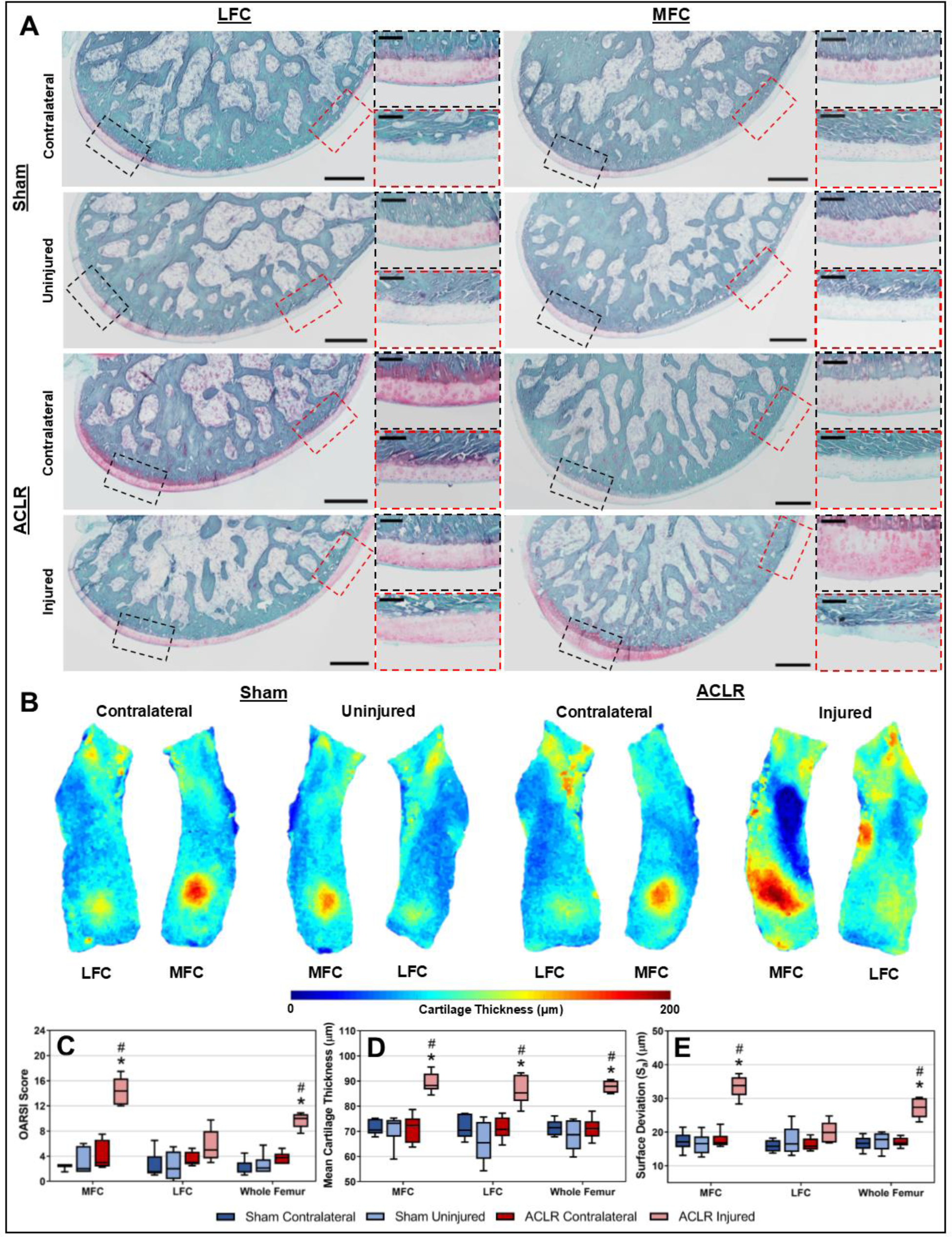
Articular Cartilage Damage and Degeneration. Histologically, the MFC and LFC of both limbs in Sham exhibit congruent articular cartilage with normal cellular morphology and staining intensity (A). Minor superficial cartilage damage was observed on the MFC of ACLR contralateral limbs. ACLR injured femora exhibited marked structural damage and hypocellularity on the anterior MFC, with swelling and abnormal chondrocyte clustering on the posterior MFC. The LFC of ACLR injured femora exhibited swelling and superficial damage. CE-μCT confirms histological observations by demonstrating drastic alterations in cartilage thickness distributions on the MFC of ACLR injured (B). ACLR injured femora had significantly increased OARSI score (n=6 Sham; n=6 ACLR) (C), increased femoral cartilage thickness (D), and increased femoral cartilage surface deviation (E). * indicates significant difference to contralateral limb; # indicates significant difference to ipsilateral limb in Sham. Scalebar of whole-condyle images is 500 μm. Scalebar of high-magnification images is 200 μm.

## DISCUSSION

This study sought to characterize acute alterations to trabecular and SCB in a rat model of noninvasive ACL rupture, and assess whether alterations in bone precede measurable cartilage changes. Our results demonstrate that in this model, rapid alterations to SCB and trabecular bone turnover and microstructure are observed immediately after injury. Despite evidence of catabolic bone remodeling as early as 3-7 days post-injury, our findings also indicate notable damage to cartilage of the MFC in ACL-injured femora by 14 days post-injury. Thus, given the magnitude of cartilage damage, we do not conclude that bony changes markedly precede AC degeneration in this model. Collectively, our results demonstrate that the acute bone loss observed after traumatic joint injury is driven by both thwarted anabolism (based on NIR BoneTag and dynamic histomorphometry results) and increased catabolism (based on NIR CatK and TRAP histology results).

Prior clinical studies have demonstrated a compartment/region-dependent loss of periarticular bone mass and/or bone mineral density following traumatic joint injury^37-42^. Several of these studies indicate incomplete recovery of bony deficits in the long-term, with some evidence even indicating lower BMD in the injured limb 10+ years after knee injury.^40^ A recent study which first employed high-resolution CT in humans with ACL injury demonstrated the microstructural nature of injury-induced bone loss^42^, largely corroborating our observations in rats. Collectively, these data point towards a phenotypic shift in periarticular bone and long-term imbalance between tissue anabolism and catabolism. The present study supports this premise, as our data demonstrate that ACLR markedly thwarts bone formation/anabolism and activates bone resorption/catabolism, leading to complex, region-dependent remodeling. Histologically, this response was characterized by decreased bone deposition and increased osteoclast numbers. While this remodeling is largely characterized by bone loss, our analysis of LFC SCB indicates a small but significant increase in SCB BV/TV and TMD. This evidence of a compartment/region-dependent bone remodeling response to injury points towards the complexity of PTOA pathogenesis and the importance of evaluating the entire joint in its individual regions when characterizing pathophysiology and evaluating therapeutic strategies.

While chronic joint destabilization is known to induce progressive cartilage degeneration, it remains unclear whether the damage observed on the MFC in this ACLR model (and similar models in mice^22^) is primarily due to chronic instability or due to acute structural damage from injury loading. Our present findings indicate a similar, albeit less severe cartilage phenotype on the MFC at this early 14-day timepoint as previously observed at both 4 and 10 weeks post-ACLR in our prior studies^26^. Interestingly, the same overall phenotype was also observed in a rat surgical ACL transection model, employed as a comparison in the same prior study^26^, which lacks injurious mechanical loading. Thus, we conclude that ACL deficiency-induced joint destabilization has an immediate impact on the onset of cartilage degeneration in this model, that this degeneration is concurrent with bony remodeling, and that it is not just the acute trauma to cartilage during the injurious subluxation and overloading event that induces the observed phenotype on the MFC. While synovitis – a non-mechanical contributor to PTOA pathogenesis – is known to also initiate cartilage degeneration via cytokines, chemokines, and tissue proteases, these effects would not be expected to be region-dependent. Thus, the drastic differences in cartilage phenotype on the anterior vs posterior MFC may be attributable to primarily mechanical factors. It is also possible that proinflammatory cytokines and proteases expressed by synovium differentially accelerate extracellular matrix damage in chondrocytes that undergo pathological loading conditions.

Our findings stress the importance of evaluating both the contralateral limb of the injured rat and a limb from a sham-loaded rat in this PTOA model. While contralateral bone and cartilage changes were subtle, they nonetheless were observable. Reduced activity leading to offloading of both limbs following joint injury, in addition to contralateral compensation and gait changes may confound interpretation of tissue changes in the injured limb if only compared to the contralateral limb. Future studies should avoid bilateral ACLR and employ appropriate controls.

This study is not without limitations. Our conclusion regarding the temporal relationship between AC degeneration and bone remodeling are based on a single endpoint assessment of AC, given the technical limitations of contrast-enhanced imaging and the destructive nature of histology. Future studies may develop *in vivo* contrast-enhanced imaging approaches to longitudinally assess AC morphology. Although our rigorous image analysis methodology sought to eliminate subjective user input via registration-based algorithms, our workflow nonetheless relied on some manual tissue contouring. Our histomorphometry and NIR data demonstrates thwarted bone formation, however we did not perform histologic osteoblast counts to determine whether this effect is driven by a reduction in osteoblast activity or overall osteoblast numbers.

In a study combining multi-disciplinary imaging, histological and histomorphometric measures to characterize acute joint injury-induced bone and cartilage remodeling, we found that noninvasive ACLR in the rat induces immediate and sustained reduction of bone anabolism and an overactivation of bone catabolism. Future studies in our group aim to elucidate how intra-articular inflammation promotes this phenotype in order to uncover the molecular mechanisms of degenerative tissue changes, facilitating the development of novel PTOA treatments.

## ACKNOWLEDGEMENTS

N/A

